# Adaptive Identification of Cortical and Subcortical Imaging Markers of Early Life Stress and Posttraumatic Stress Disorder

**DOI:** 10.1101/482448

**Authors:** Lauren E. Salminen, Rajendra A. Morey, Brandalyn C. Riedel, Neda Jahanshad, Emily L. Dennis, Paul M. Thompson

**Affiliations:** Imaging Genetics Center, Mark and Mary Stevens Institute for Neuroimaging & Informatics, Keck School of Medicine, University of Southern California, 4676 Admiralty Way, Marina del Rey, CA 90292; Durham VA Medical Center, Durham, NC; Duke University Medical Center, Durham, NC; Department of Radiology and Imaging Sciences, Indiana University School of Medicine, Indianapolis, IN; Psychiatry Neuroimaging Laboratory, Harvard Medical School, Boston, MA; Stanford Neurodevelopment, Affect, and Psychopathology Laboratory, Stanford, CA

**Author notes:** Corresponding Author contact information, 323-44-BRAIN.

**Keywords:** PTSD, early life stress, neuroimaging, machine learning

## Abstract

**Background and Purpose:** Posttraumatic stress disorder (PTSD) is a heterogeneous condition associated with a range of brain imaging abnormalities. Early life stress (ELS) contributes to this heterogeneity, but we do not know how a history of ELS influences traditionally defined brain signatures of PTSD. Here we used a novel machine learning method - evolving partitions to improve classification (EPIC) - to identify shared and unique structural neuroimaging markers of ELS and PTSD in 97 combat-exposed military veterans.

**Methods:** We used EPIC with repeated cross-validation to determine how combinations of cortical thickness, surface area, and subcortical brain volumes could contribute to classification of PTSD (n=40) versus controls (n=57), and classification of ELS within the PTSD (ELS^+^ n=16; ELS^-^n=24) and control groups (ELS^+^ n=16; ELS^-^ n=41). Additional inputs included intracranial volume, age, sex, adult trauma, and depression.

**Results:** On average, EPIC classified PTSD with 69% accuracy (SD=5%), and ELS with 64% accuracy in the PTSD group (SD=10%), and 62% accuracy in controls (SD=6%). EPIC selected unique sets of individual features that classified each group with 75-85% accuracy in post hoc analyses; combinations of regions marginally improved classification from the individual atlas-defined brain regions. Across analyses, surface area in the right posterior cingulate was the only variable that was repeatedly selected as an important feature for classification of PTSD and ELS.

**Conclusions:** EPIC revealed unique patterns of features that distinguished PTSD and ELS in this sample of combat-exposed military veterans, which may represent distinct biotypes of stress-related neuropathology.

## Introduction

PTSD is a serious mental health condition in which patients often show brain abnormalities in regions involved in memory, fear, and emotional processing - namely the hippocampus and amygdala - and regions of the prefrontal and cingulate cortex.^1^ However, the etiology of PTSD is complicated by individual differences in predisposing factors that may also affect brain structure and symptoms.^2-3^ There is interest in identifying disease mechanisms and risk factors that influence brain outcomes among individuals with PTSD (PTSD^+^), and early life stress (ELS) is one key predisposing factor that affects brain abnormalities in this population.^4-5^

ELS affects cortical and subcortical brain structures implicated in PTSD,^6^ but studies show that ELS and PTSD do not impact brain structure identically.^7-8^ In a study of military veterans,^9^ PTSD symptom severity was related to cortical thickness (CT) in the posterior cingulate (PCC)/paracentral area, but the direction of this association depended on ELS. Specifically, CT in these regions was positively associated with PTSD symptom severity in ELS-exposed (ELS^+^) veterans, but negatively associated with symptom severity in ELS-unexposed (ELS^-^) veterans. Further, associations of symptom severity with amygdala and hippocampal volumes were significant only in the ELS^+^ group. ELS may therefore have a regionally selective influence on the brain that leads to different profiles of abnormalities in trauma exposed veterans with and without PTSD. We do not fully understand how PTSD affects cortical surface area (SA), but lower SA has been reported in maltreated children and may have implications for PTSD^+^ patients with a history of ELS.^10^

We recently developed a new supervised machine learning tool, evolving partitions to improve classification (EPIC), that adaptively selects and merges sets of brain measures to improve group classification.^11^ EPIC is based on the idea that certain features may not relate to a grouping variable when examined independently, but when integrated with others, may better discriminate subjects from different classes. This top-down approach reduces the dimension of a set of imaging predictors by re-partitioning the cortex into regions and “super-regions” (the result of merging 2 or more regions) to boost group classification. EPIC was initially used to denote “evolving partitions to improve connectomics” -^11^ it was used to cluster and simplify connectivity matrices to achieve better classification. In general, the adaptive merging process can be applied to improve classification based on clusters of any type of spatial data, including the structural data used here, so here we use the term to refer to “evolving partitions to improve classification”. Our group previously showed that EPIC improves the accuracy of disease classification in Alzheimer’s disease^11^ and traumatic brain injury.^12^ Here we used EPIC to determine if unique combinations of SA, CT, and subcortical volumes (VL) could discriminate individuals with PTSD and ELS. Thus, our primary goal in this study was to identify new patterns of neuroimaging signatures that may represent unique phenotypes of PTSD or ELS and could be tested in subsequent studies in both military and civilian samples.

## Method

### Participants

Data were collected from 97 combat-exposed Operation Enduring Freedom and Operation Iraqi Freedom (OEF/OIF) veterans (PTSD n=40, controls n=57) who took part in the military conflicts in Iraq and Afghanistan following the September 11^th^ 2001 terrorist attacks. Participants were aged 23-65 (male, n=80; female, n=17) and recruited from the Durham VA and Duke University Medical Centers. Exclusion criteria consisted of any Axis I diagnosis other than PTSD or major depressive disorder, substance dependence (other than nicotine), high risk for suicide, history of learning disability or developmental delay, history of head injury with loss of consciousness > 5 min, neurological disorders, major medical conditions, and contraindications for MRI (e.g., claustrophobia). The same exclusion criteria applied to controls, except that control subjects could not meet criteria for any Axis I diagnosis. We did not exclude individuals who were taking medication for depression, anxiety, and/or sleep disturbances as these conditions are prevalent in this population. All participants provided informed consent; procedures were approved by the local IRBs.

### Clinical Assessment

A diagnosis of PTSD was determined using the Clinician Administered PTSD scale for DSM-IV. Adult trauma and ELS exposure were evaluated using the Traumatic Life Events Questionnaire (TLEQ).^13^ Adult trauma was quantified as a continuous variable reflecting exposure severity (i.e., total number of exposures). Participants indicating exposure to any traumatic event before age 18 were identified as ELS^+^, whereas participants indicating no trauma exposure before age 18 were identified as ELS^-^ (unexposed).^14^ Current depression was measured using the Beck Depression Inventory-II (BDI).^15^

### Neuroimaging Acquisition and Processing

MRI scanning was completed at two sites using a 3T GE MR750 (93% of sample) and 3T GE Signa EXCITE (7% of sample) scanner, each equipped with an 8-channel head coil. Chi-squared analyses revealed no significant differences between sites and target groups (PTSD by site, *x*^2^(1)=0.79, p=0.375; ELS by site, *x*^2^(1)=0.33, p=0.564). Independent of the target groups, we did not observe site differences in the distribution of sex (*x*^2^(1)=1.6, p=0.205), age (t(95)= -0.86, p=0.393), BDI scores (t(95)= -1.24, p=0.216), or adult trauma exposure (t(95)= -0.33, p=0.743). T1-weighted axial brain images were obtained with 1-mm isotropic voxels using the following parameters: GE Signa EXCITE: TR/TE/flip=8.208-ms/3.22-ms/12°, GE MR750: TR/TE/flip=7.484-ms/2.984-ms/12°; all images were captured with FOV=256 × 256mm, 1-mm slice thickness. T1-weighted images were processed using FreeSurfer version 5.3. Cortical and subcortical extractions were performed and checked for quality control using standardized protocols from the ENIGMA consortium, yielding CT and SA for 34 bilateral cortical regions of interest (ROIs), and bilateral VL for 8 subcortical structures.

### Design

Primary analyses were completed to determine if EPIC could distinguish PTSD^+^ veterans from trauma-exposed controls using 152 neuroimaging inputs (68 CT, 68 SA, and 16 VL). EPIC was also used to determine the role of ELS within each group (PTSD: N=40, ELS^+^ (n=16) /ELS^-^ (n=24); Controls: N=57, ELS^+^ (n=16) /ELS^-^ (n=41)). All analyses included age, sex, and intracranial volume (ICV) as input variables. Preliminary analyses revealed no significant differences in age, sex, BDI scores, TLEQ scores, PTSD, and ELS by site, therefore we did not include site/scanner as input features. Total score on the BDI was included as an input feature when depression differed significantly between the target groups in order to model depression symptoms as an important feature of PTSD and ELS. We further tested whether EPIC or the original FreeSurfer segmentation could distinguish target groups after regressing out the effects of depression on the input features when BDI scores differed significantly between groups. We hypothesized that both classifiers would perform worse when the BDI was removed as input feature for analyses where depression differed between groups.

### Classification with EPIC

The workflow of EPIC is shown in Figure 1. Here input variables included CT, SA, and VL measurements, and the respective covariates for each analysis. Structural MRI inputs were used to generate merged composite cortical partitions through iterative ROI sampling over the space of possible ROI groupings. We applied a 70/30 (training/testing) stratified split of the data, in which a linear support vector machine (SVM) learned the optimal partitions for group classification in the training data, and then evaluated the partitions in the test data using stratified *k*-fold cross-validation (CV). After splitting the data, 67 participants were used for *k*-fold CV and 30 participants were held out for testing. In line with original group distributions, we maintained 40% cases and 60% controls for each analysis, resulting in 12 cases/18 controls for hold-out, and 27 cases/40 controls for training. For the primary analysis of PTSD vs. controls, we applied 5-fold CV to the training data, resulting in approximately 5 PTSD cases and 8 controls per fold. For secondary analyses of ELS within the PTSD and control groups, we used 2-fold CV to account for lower cell sizes in the target subgroups. We then re-based the stratified CV rates according to 40% of ELS^+^ participants and 60% of ELS^-^ participants within each subgroup. For the analysis of ELS in PTSD patients, the re-based CV rate provided a total test set of 6 ELS^+^ and 14 ELS^-^ PTSD patients, divided by 2 (folds), equaling 3 ELS^+^ and 7 ELS^-^ patients per fold. For the analysis of ELS in controls, the re-based cross-validation rate provided a total test set of 6 ELS^+^ and 25 ELS^-^ controls, divided by 2 (folds), equaling 3 ELS^+^ and 13 ELS^-^ controls per fold. *K*-fold cross validation was repeated 10 times for all analyses. Simulated annealing was used to better identify optimal partitions, or ways of merging structural measures.^11^ The EPIC algorithm was written in Python using an in-house modification of scikit learn.^16^ The code is available from the authors upon request.

**Figure 1.**
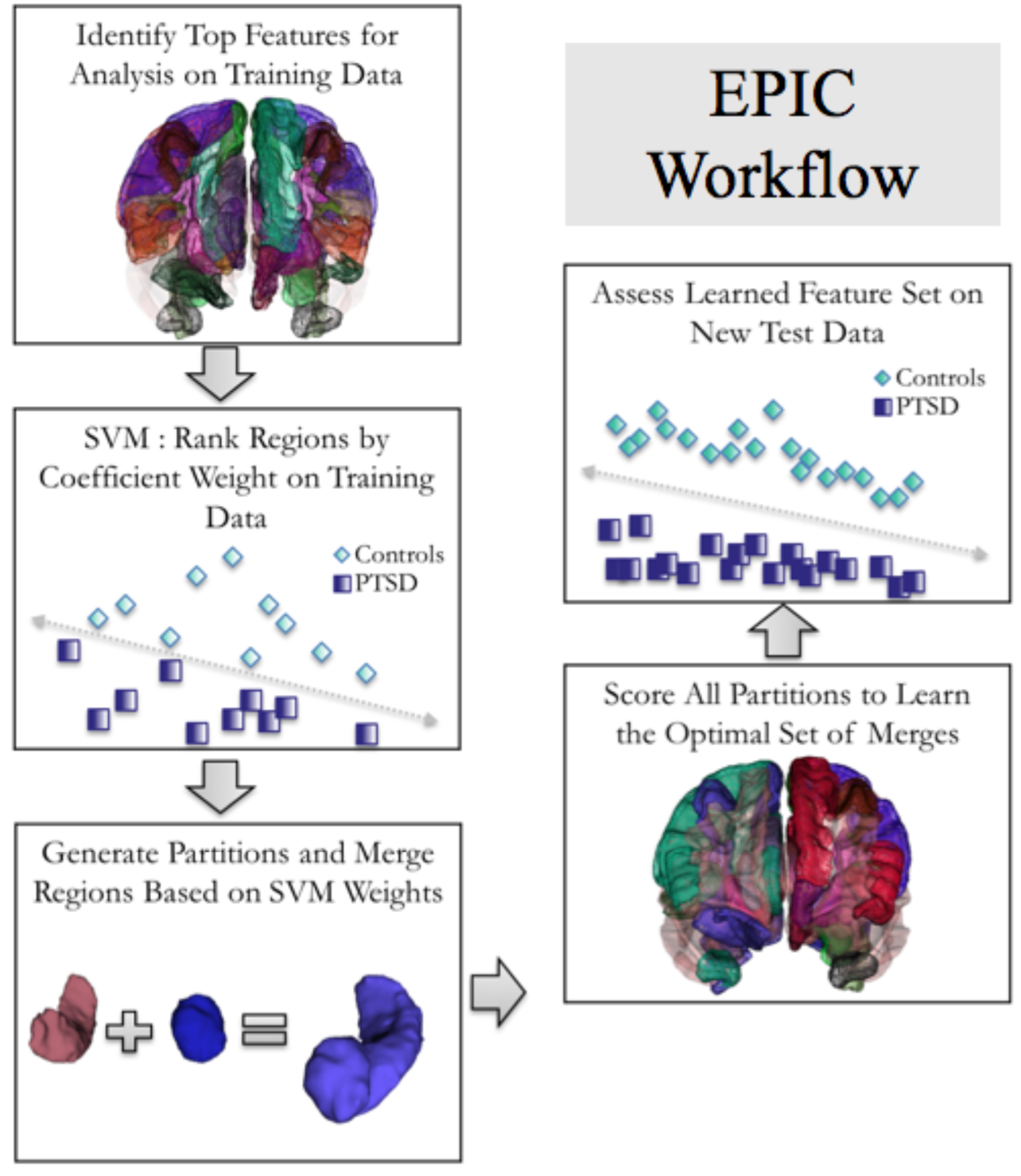
Diagram of the combinatorial support vector machine (SVM) approach used by EPIC. Labels depict the partitions used for classifying PTSD from trauma-exposed controls in the main analysis.

We evaluated EPIC’s performance using balanced accuracy ([(true positive (TP)/(TP + false negative (FN)) + true negative (TN)/(TN + false positive (FP))]/2), sensitivity (TP/TP+FN), specificity (TN/FP +TN), and the positive predictive value (PPV; TP/TP+FP) averaged across the 10 repeats. We also report performance metrics for the top repeats of EPIC, which we defined as classification accuracy ≥ 65%, to demonstrate the importance of repeated CV in machine learning studies. Balanced accuracy was used as our primary index of performance to account for modest sample size imbalances between the target groups. We include three additional metrics of classifier performance for each analysis in Table 1, to allow for comparison to others that have been reported in the literature. To determine whether EPIC’s combinatorial approach improved classification of the target groups, we report classifier performance for the SVM using the original FreeSurfer brain measures and covariates modeled individually. The same optimization parameters were applied to both sets of models. Using a thresholding feature elimination procedure, individual regions and super-regions with absolute SVM coefficient weights (*w*) <0.05 on training data were considered noise and removed from the primary analysis to enhance power. We set a threshold of 0.003 for the secondary analyses, which was determined from the final accuracy outputs from the test data.

**Table 1.** Average Performance Results Across 10 Repeats

| Table 1. Average Performance Results Across 10 Repeats |  |  |  |  |  |  |  |
| --- | --- | --- | --- | --- | --- | --- | --- |
| Classification with features modeled individually |  |  |  |  |  |  |  |
| Model | Acc | bAcc* | Sens* | Spec* | PPV* | F1 | G<sub>mean</sub> |
| PTSD vs. Controls | 68% | 67% | 56% | 77% | 65% | 0.60 | 66% |
| ELS<sup>+</sup> vs. ELS<sup>-</sup>, PTSD | 64% | 63% | 56% | 70% | 58% | 0.54 | 63% |
| ELS<sup>+</sup> vs. ELS<sup>-</sup>, Controls | 68% | 57% | 30% | 84% | 42% | 0.33 | 50% |
| Classification with EPIC |  |  |  |  |  |  |  |
| Model | Acc | bAcc* | Sens* | Spec* | PPV* | F1 | G<sub>mean</sub> |
| PTSD vs. Controls | 71% | 69% | 58% | 81% | 69% | 0.62 | 69% |
| ELS<sup>+</sup> vs. ELS<sup>-</sup>, PTSD | 66% | 64% | 53% | 75% | 60% | 0.54 | 63% |
| ELS<sup>+</sup> vs. ELS<sup>-</sup>, Controls | 70% | 62% | 43% | 81% | 48% | 0.44 | 59% |

*Abbreviations and formulas:* Acc= Accuracy, true positive (TP)+true negative (TN)/TP+TN+false positive (FP)+false negative (FN); bAcc = balanced accuracy, [(TP/(TP+FN)+TN/(TN+FP))/2]; Sens= sensitivity (recall), TP rate, TP/TP+FN; Spec= specificity, TN rate, TN/FP +TN; PPV= positive predictive value (precision), TP/TP+FP; F1= harmonic mean of precision and recall, $2 \times (\text{Recall} \times \text{Precision}) / (\text{Recall} + \text{Precision})$ ; G<sub>mean</sub>= geometric mean, $\sqrt{\text{TPrate} \times \text{TNrate}}$ ; \*reported in main text.

## Results

### PTSD vs. trauma-exposed controls

PTSD^+^ veterans scored higher (*M*=15.9, *SD*= 11.2) on the BDI compared to controls (*M*=4.2, *SD*=5.4, *p*<0.001), so the total BDI score was included as an input feature. EPIC classified PTSD with 69% accuracy (SD=5%) on average, with 58% sensitivity, 81% specificity, and 69% PPV. Best performance with EPIC classified PTSD from trauma-exposed controls with 77% accuracy (63% sensitivity, 91% specificity, 83% PPV, Figure 2C); seven out of 10 repeats classified PTSD with greater than 65% accuracy. Across these repeats, EPIC identified the BDI as the strongest predictor of PTSD compared to all imaging variables. Super-regions were identified as important features across 5 out of the 7 top repeats but did not contribute to accuracy for the best individual repeat with EPIC (Figure 2). In Figure 2H, we report the most frequently selected features across the top repeats of EPIC. After the BDI, the strongest predictors of PTSD were CT in the left temporal pole (TP), and SA in the left post central gyrus, and left inferior temporal gyrus (ITG). Compared to average performance using the individual input features, EPIC modestly improved classification accuracy by 2%, on average (Table 1).

**Figure 2.**
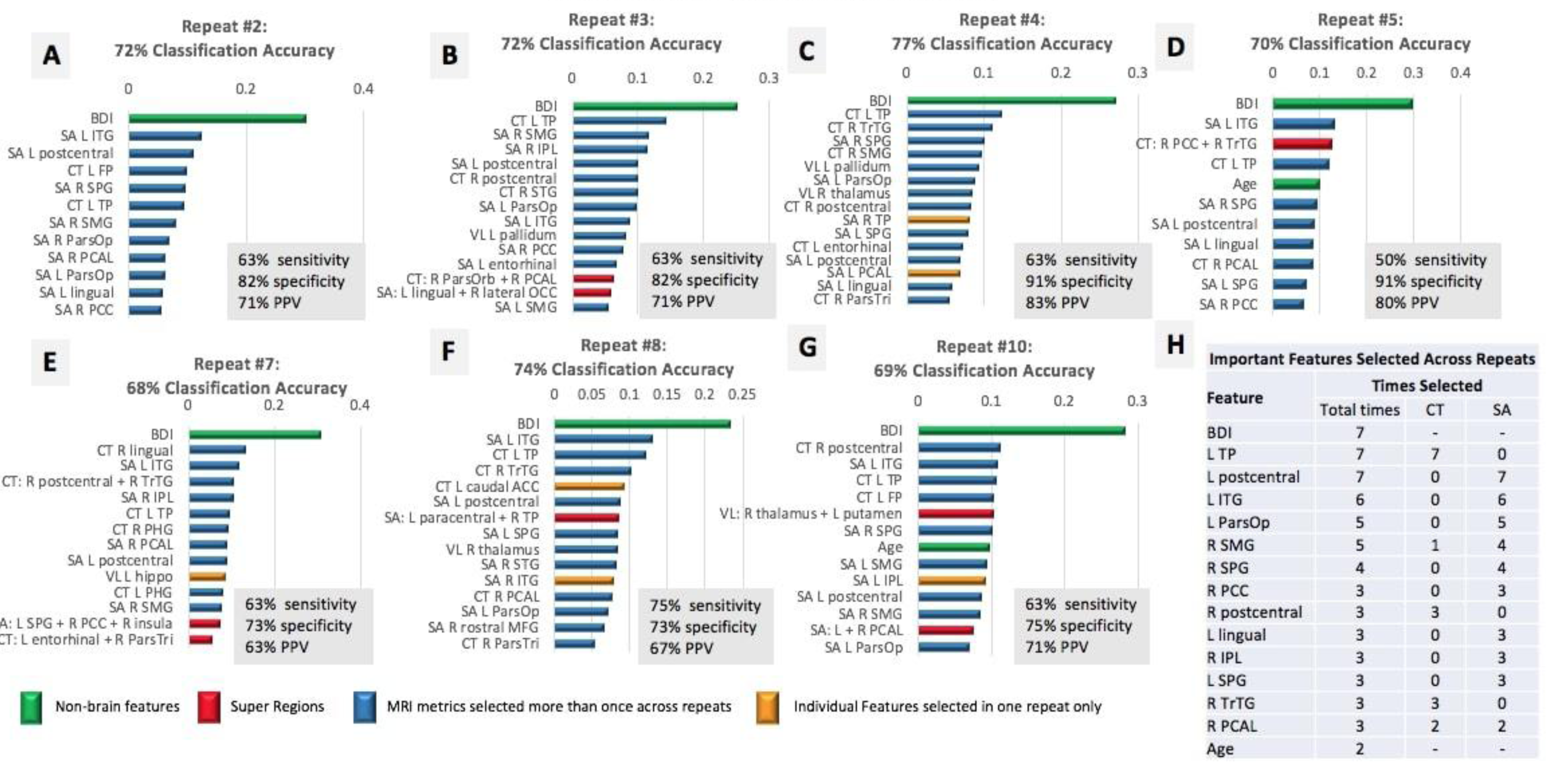
Panels A-G. The top features for the strongest repeats of EPIC (≥65% classification accuracy) are plotted in rank order according the absolute value of the SVM coefficient. We limited the number of features shown to the top 15 in each feature list, but most repeats revealed less than 15 features that contributed to final classification accuracy. The most important features are reported (**Panel H**) in rank order according to the frequency by which each variable was identified as a top predictor of PTSD across the top repeats of EPIC. Variables that were selected the same number of times across repeats were ranked by their relative position and coefficient weight across feature lists of each individual repeat. Most MRI features were selected more than once across repeats (*blue*); features selected in one repeat are displayed in *yellow*. Super regions (*red*) consisted of 3 or less merged regions in this analysis. Non-brain regions (*green*) were selected in each repeat, with the BDI selected as the most important feature in each repeat. **Acronyms** (in descending order from left to right): SA (surface area), CT (cortical thickness), VL (subcortical volumes), L (left hemisphere), R (right hemisphere), BDI (score on the Beck Depression Inventory), ITG (inferior temporal gyrus), FP (frontal pole), SPG (superior parietal gyrus), TP (temporal pole), SMG (supramarginal gyrus), ParsOp (pars opercularis), PCAL (pericalcarine), PCC (posterior cingulate), IPL (inferior parietal lobule), STG (superior temporal gyrus), ParsOrb (pars orbitalis), ParsTri (pars triangularis), PHG (parahippocampal gyrus), hippo (hippocampus), TrTG (transverse temporal gyrus), ACC (anterior cingulate), MFG (middle frontal gyrus).

Model performance declined significantly when we tested the classifier on inputs that were residualized for BDI scores. Neither the original FreeSurfer segmentation, nor the EPIC classifier could distinguish PTSD groups from controls, with the same low discriminatory power between methods (38% accuracy, on average). Given this low accuracy, we did not apply this approach for other analyses that differed on BDI scores.

### PTSD^+^ veterans with and without ELS

PTSD^+^ groups did not differ significantly on BDI scores (ELS^+^: *M*=16.4, *SD*=12.2; ELS^-^: *M*=15.5, *SD*=10.7) or adult trauma exposure (ELS^+^: *M*=13.6, *SD*=8.5; ELS^-^: *M*=10, *SD*=10.1) indexed with the TLEQ, so these variables were not used here as input features. On average, EPIC classified ELS in the PTSD group with 64% accuracy (SD=10%), 56% sensitivity, 70% specificity, and 60% PPV. Best performance with EPIC classified ELS with 77% accuracy (63% sensitivity, 92% specificity, 83% PPV Figure 3C); five of the ten repeats classified ELS with greater than 65% accuracy. EPIC repeatedly selected VL in the right putamen, and CT in the left caudal anterior cingulate (ACC) and right parahippocampal gyrus (PHG) as the strongest predictors of ELS (Figure 3F). Super regions were not selected as important features for distinguishing ELS across the top repeats. In comparison to the SVM that modeled features individually, EPIC marginally improved classification accuracy by 2% (Table 1).

**Figure 3.**
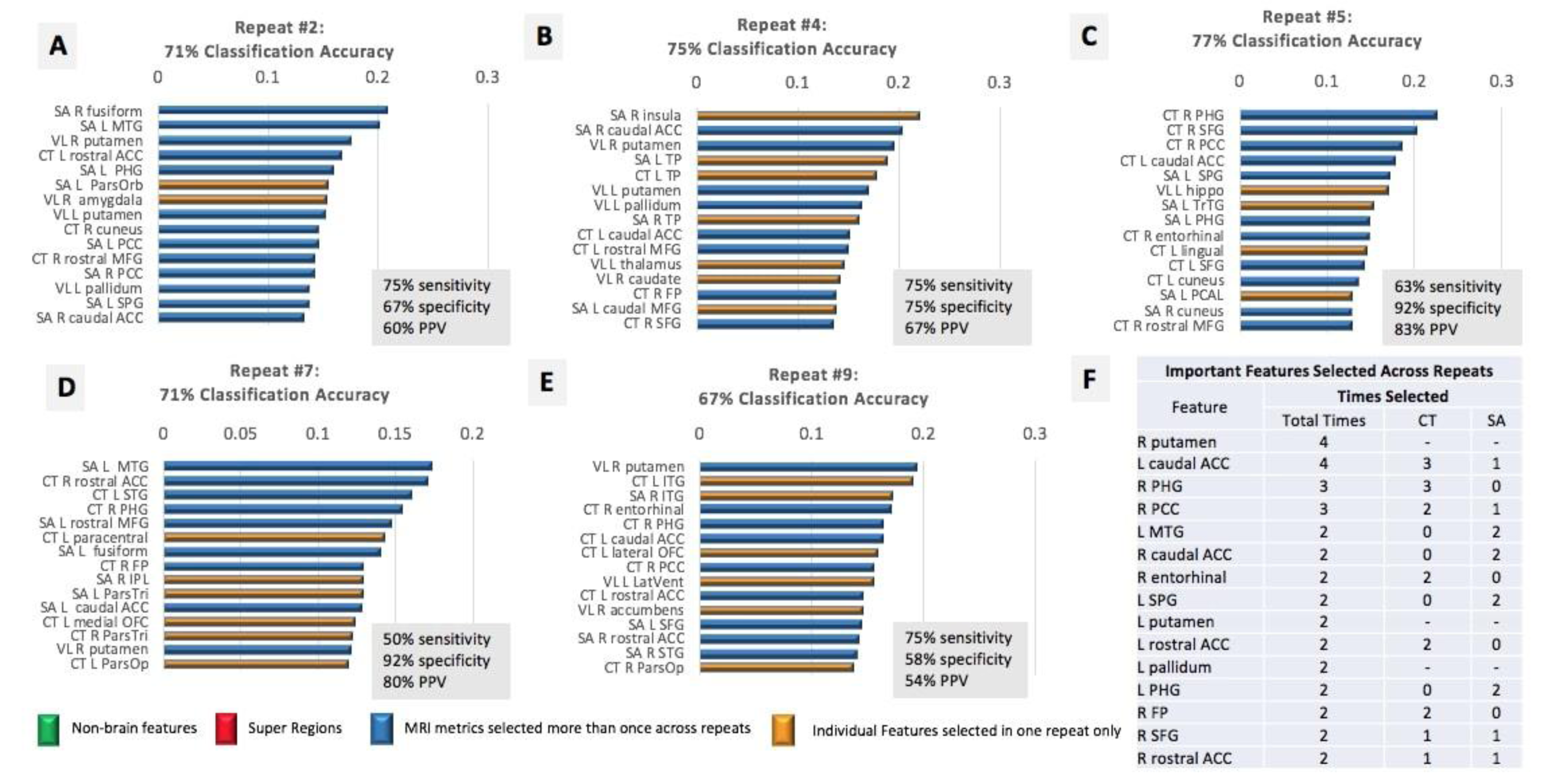
Panels A-E. The top features for the strongest repeats of EPIC (≥65% classification accuracy) are plotted in rank order according the absolute value of the SVM coefficient. We limited the number of features shown to the top 15 in each feature list. The most important features selected across the top repeats of EPIC are reported in rank order (**Panel F**) according to the frequency by which each variable was identified as a top predictor of ELS in the PTSD group. Variables that were selected the same number of times across repeats were ranked by their relative position and coefficient weight across feature lists. Most MRI features were selected more than once across repeats (*blue*); features selected in one repeat are displayed in *yellow*. Non-brain metrics (*green*) were not important for ELS classification in this analysis. Although super regions were identified in the complete feature list of each repeat, they were among the weakest predictors and not plotted here. Non-brain regions (*green*) were selected in each repeat, with the BDI selected as the most important feature in each repeat. **Acronyms** (in descending order from left to right): SA (surface area), CT (cortical thickness), VL (subcortical volumes), L (left hemisphere), R (right hemisphere), MTG (middle temporal gyrus), ACC (anterior cingulate), PHG (parahippocampal), ParsOrb (pars orbitalis), PCC (posterior cingulate), MFG (middle frontal gyrus), SPG (superior parietal gyrus), TP (temporal pole), FP (frontal pole), SFG (superior frontal gyrus), hippo (hippocampus), TrTG (transverse temporal gyrus), PCAL (pericalcarine), STG (superior temporal gyrus), IPL (inferior parietal lobule), ParsTri (pars triangularis), ParsOp (pars opercularis), OFC (orbitofrontal cortex), LatVent (lateral ventricle).

### Trauma-exposed controls with and without ELS

BDI scores and adult trauma exposure differed significantly between ELS^+^ (BDI, *M*=6.9, *SD*=7.1; TLEQ, *M*=13.1, *SD*=5.8) and ELS^-^ controls (BDI, *M*=3.1, *SD*=4.2, *p*=0.016; TLEQ, *M*=13.1, *SD*=5.8, *p*=0.001), so these variables were included as covariates in this analysis. On average, EPIC classified ELS with 62% accuracy (SD=6%), 43% sensitivity, 81% specificity, and 48% PPV. Three out of 10 repeats achieved accuracy ≥ 65%, and the best performance with EPIC classified participants with ELS with 71% accuracy (63% sensitivity, 80% specificity, 67% PPV; Figure 4A). One super-region of CT was identified as a top feature in the 9^th^ repeat of EPIC (Figure 4B), but sensitivity and PPV here were low (50%). Adult trauma indexed on the TLEQ was the only input feature that was repeatedly selected as an important predictor of ELS across the top three repeats. Several features were selected in two of the three repeats (Figure 4D), including CT in the right isthmus cingulate (ICC) and pericalcarine, SA in the right supramarginal gyrus (SMG) and posterior cingulate (PCC), and total scores on the BDI. EPIC improved classification accuracy from the original FreeSurfer segmentation by 4% (Table 1).

**Figure 4.**
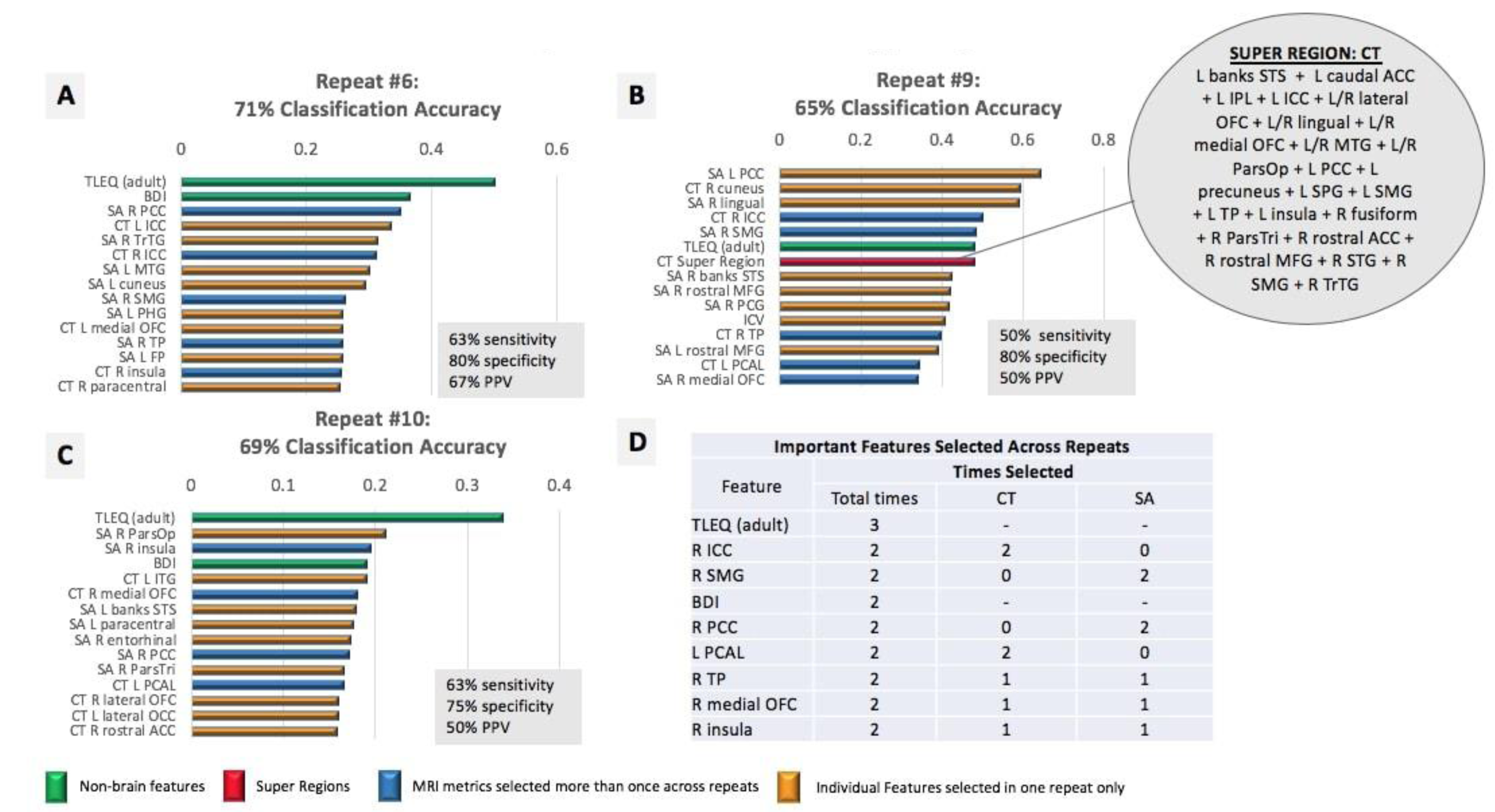
Panels A-C. The top features for the strongest repeats of EPIC (≥ 65% classification accuracy) are plotted in rank order according the absolute value of the SVM coefficient. Only three repeats achieved accuracy ≥ 65%. We limited the number of shown features to the top 15 in each repeat. The most important features selected across the top repeats of EPIC are reported in rank order (**Panel D**), according to the frequency by which each variable was identified as a top predictor of ELS in the control group. Variables that were selected the same number of times across repeats were ranked by their relative position and coefficient weight across feature lists. Most MRI features were selected once across repeats (*yellow*); features selected more than once are displayed in *blue.* Non-brain metrics (*green*) were identified as important features in each repeat, with adult trauma exposure collectively selected as the most important feature for ELS classification in this analysis. One super region (*red*) was identified as a top feature in the 9^th^ repeat of EPIC (**Panel B**). It consisted of merged regions of cortical thickness (CT). Acronyms (in descending order from left to right): SA (surface area), L (left hemisphere), R (right hemisphere), TLEQ (score on the Traumatic Life Events Questionnaire-adult), BDI (score on the Beck Depression Inventory), PCC (posterior cingulate), ICC (isthmus cingulate), MTG (middle temporal gyrus), SMG (supramarginal), PHG (parahippocampal gyrus), OFC (orbitofrontal cortex), TP (temporal pole), FP (frontal pole), banks STS (banks of superior temporal sulcus), MFG (middle frontal gyrus), ICV (intracranial volume), PCAL (pericalcarine), ParsOp (pars opercularis), ITG (inferior temporal gyrus), ParsTri (pars triangularis), lateral OCC (lateral occipital cortex), ACC (anterior cingulate).

### Observed feature patterns across the analyses

In Table 2 we provide a ranked list of unique and shared features of the target groups based on the repeated selection of variables that were among the top discriminating predictors in each analysis. SA in the right PCC was the only feature that was repeatedly selected as an important predictor for each analysis. As a post-hoc analysis, we used logistic regression to test whether the unique feature patterns identified in each analysis would explain a significant portion of variance between the target groups. The PCC was included as an independent variable in all three analyses. For the main analysis, EPIC’s feature pattern classified participants with PTSD at 74% accuracy (Nagelkerke R^2^=0.36); the strongest individual predictors were greater SA in the left *pars opercularis* (p=0.007, β=0.89) and left lingual gyrus (p=0.002, β=1.2). In the PTSD group, EPIC’s feature pattern classified participants with ELS at 85% accuracy (Nagelkerke R^2^=0.78). No individual predictors were significant at the 0.05 alpha level, but lower CT in the rostral ACC (p=0.056, β=-3.4) and PCC (p=0.069, β=-4.4), and *greater* CT in the caudal ACC (p=0.062, β=4.03) trending towards significance. Finally, the observed feature pattern in controls classified participants with ELS at 83% accuracy (Nagelkerke R^2^=0.57), with greater adult trauma exposure (p=0.007, β=1.7) and lower CT in the right medial OFC (p=0.015, β=-1.5) as the strongest predictors in the model.

**Table 2.** Unique and Shared Input Features by Analysis

| Pattern 1:<br>PTSD | Pattern 2:<br>ELS+PTSD | Pattern 3:<br>ELS | Shared features |
| --- | --- | --- | --- |
| L postcentral (SA) | R putamen (VL) | TLEQ (adult) | R PCC (SA)* |
| L ITG (SA) | L caudal ACC (CT/SA) | R ICC (CT) |  |
| L ParsOp (SA) | R PHG (CT) | R medial OFC (CT/SA) |  |
| R postcentral (CT) | L MTG (SA) | R insula (CT/SA) |  |
| L lingual (SA) | R caudal ACC (SA) |  |  |
| R IPL (SA) | R entorhinal (CT) |  |  |
| R TrTG (CT) | L putamen (VL) |  |  |
| Age | L rostral ACC (CT) |  |  |
|  | L pallidum (VL) |  |  |
|  | L PHG (SA) |  |  |
|  | R FP (CT) |  |  |
|  | R SFG (CT/SA) |  |  |
|  | R rostral ACC (CT/SA) |  |  |

*Note.* Variables in the first three columns are features that were repeatedly selected as important predictors of the target group of the respective column, and were not repeatedly selected as an important feature in other analyses. The last column indicates variables selected as important features across the analyses. **Acronyms** (in descending order from left to right): L (left hemisphere), R (right hemisphere), SA (surface area), ITG (inferior temporal gyrus), ParsOp (pars opercularis), CT (cortical thickness), PCC (posterior cingulate), IPL (inferior parietal lobule), TrTG (transverse temporal gyrus), VL (subcortical volumes), ACC (anterior cingulate), PHG (parahippocampal gyrus), MTG (middle temporal gyrus), FP (frontal pole), SFG (superior frontal gyrus), TLEQ (traumatic life events questionnaire), ICC (isthmus cingulate), OFC (orbitofrontal cortex).

## Discussion

Our machine learning method, EPIC, revealed preliminary neuroimaging signatures of PTSD and ELS that classified participants with greater than 60% accuracy, on average, across analyses. BDI scores were identified as an important feature for distinguishing PTSD in the whole sample and ELS in trauma exposed controls, suggesting a critical role of trauma-related emotional dysregulation that occurs independent of a PTSD diagnosis. Comparison of the learned features that were repeatedly selected in each analysis revealed three unique patterns of neuroimaging features, with common involvement of the PCC – an important brain region for processing emotionally salient stimuli.^17-18^ These patterns may represent underlying “biotypes” of childhood and adult trauma, which we discuss in detail below.

Classification was strongest when distinguishing PTSD from trauma-exposed controls (69% on average), with total scores on the BDI repeatedly selected as the most important feature across all repeats of EPIC. The most important neuroimaging features included metrics of cortical regions involved in somatosensory function, social cognition, and emotional processing of sensory input. These regions distinguished individuals with PTSD *after* total BDI scores, suggesting that traditionally defined PTSD brain abnormalities in the hippocampus and amygdala may be due to co-morbid depression rather than features unique to PTSD (e.g., hypervigilance). Features that were unique to PTSD have been described as constituents of the extra-striate ventral visual cortex.^19^ When we examined the relative importance of these features as predictors of PTSD, we found larger SA in the lingual gyrus and *pars opercularis*– regions that have shown abnormal imaging results in relation to trauma and psychosis-spectrum symptoms across psychiatric diagnoses.^20-22^ The metric-specificity of these results may implicate a developmental risk for PTSD - SA expands throughout childhood and adolescence and thus is vulnerable to early environmental influence.^23^ Recent work using resting state fMRI shows distinct functional specialization of cortical regions within the extra-striate ventral visual cortex in human newborns, including unique functional associations between the cortical regions that repeatedly classified PTSD from controls.^19^Replication efforts are needed to determine the reliability of a cortico-limbic-somatosensory imaging pattern in relation to PTSD and its specific symptom clusters. Twin studies will be helpful for elucidating a potential risk for PTSD that may be tied to this learned neural profile.

The top predictors of ELS in the PTSD^+^ group included brain regions that tap emotion regulation, reward sensitivity, and executive control – functional domains that tend to be abnormal in PTSD.^24^ Among these regions was the rostral and caudal ACC, a structure that is commonly disrupted in ELS^+^ populations.^6^ ELS may prime brain regions that subserve these functions to exhibit a chronic and exaggerated threat response that disrupts brain structure and increases risk for PTSD following adult trauma exposure. The interpretation of directional differences between the caudal and rostral ACC in our post hoc analysis is unclear but may reflect a nuisance result of the multivariate design. This is an interesting topic for future work given the large body of literature implicating the ACC in stress-related phenotypes.

Adult trauma exposure was the most important feature for classifying ELS in trauma-exposed controls. This is consistent with evidence that ELS is linked with high risk for subsequent trauma exposure in adults,^25^which may be due to increased risk-taking behaviors in ELS^+^individuals.^26^Additional work shows significantly higher prevalence of ELS in military samples, which may reflect an escape from adversity among those who voluntarily enlist.^27^ The observation that depression was an important predictor of ELS also is consistent with studies that show higher symptoms of depression and emotional dysregulation among otherwise healthy individuals with a history of ELS compared to unexposed controls.^28-29^ The unique neuroimaging pattern that distinguished ELS involved the ICC, medial OFC, and insula, with the strongest post hoc predictive value in the medial OFC. This cluster of regions has been previously associated with negative affect and self-referential processing,^30^ which is consistent with a higher degree of emotional dysregulation observed in the ELS^+^ group.

Several limitations should be acknowledged. 1) EPIC combined neuroimaging features to boost classification accuracy in each analysis, but super-regions showed low feature importance relative to the individual regions. Combining neuroimaging metrics into super-regions may reduce noise in the individual measures that are most relevant to group classification, thereby improving accuracy relative to using each feature individually.^11^ 2) Sensitivity was low in all analyses, ranging from 43-58%, on average. This is not necessarily surprising because all participants in this study were exposed to military combat. Lifetime trauma exposure in military veterans may yield a unique phenotype that is independent of PTSD and ELS, limiting the detection rate of either “condition”. Sensitivity was lowest for detection of ELS in the control group (43% accuracy, on average), likely because the control group consists of non-clinical participants who should not have any gross brain abnormalities. Of note, PPVs were better than sensitivity outcomes in the PTSD analyses, and slightly better in the analysis of controls. This indicates a higher probability that individuals who screen positive for the target group truly belong to that group. The high specificity observed in controls with ELS also suggests that EPIC can distinguish participants with subtle brain differences that are within the normal range of variance that would be expected in a non-clinical sample. 3) Although our sample size is consistent with several previous studies of machine learning in neuroimaging and psychiatry, numbers were small when considering the subdivisions of cases and controls used for training and testing; this may have limited our ability to detect more robust effects across the target groups. However, recent work from the ENIGMA consortium reveals comparable rates of classification accuracy for bipolar disease in over 3,000 individuals,^31^ suggesting that sample size was not an issue in our study. Using the Research Domain Criteria (RDoC) approach to guide future machine learning studies may significantly improve classification and prediction of complex psychiatric diseases;^32^ this is currently a goal of several Working Groups within the ENIGMA consortium. 4) We did not examine the ability of EPIC to distinguish ELS^+^ from ELS^-^ controls independent of PTSD because there are multiple projects within the ENIGMA consortium that are pursuing this research topic in larger samples with complex psychiatric histories.^33^ Thus, we cannot determine from the information presented whether PTSD is easier to detect and classify than ELS. This is an important research question to address using clinically diverse datasets to provide a more powerful depiction of ELS phenotypes independent of psychiatric illness. 5) This cohort was exposed to military combat and results may not generalize to civilians. However, the most salient predictive features of PTSD and ELS are consistent with civilian studies, so they may represent the larger PTSD^+^ and ELS^+^ populations.

EPIC adapts regions of interest to improve classification, and a similar approach could be implemented for functional imaging data. Here we focused on structural MRI features to compare classifier results to the large body of literature showing structural brain disruptions among individuals with PTSD and ELS. In functional connectivity analyses, however, the seed regions that act as nodes of the network could be adaptively refined to improve classification of the target groups, and some regions could be merged or split to adapt the set of predictors. This is a current goal of the PGC-ENIGMA PTSD Working Group - the larger data source from which this work stems.

In sum, our results show region-specific distinctions in the neuroimaging profiles of PTSD and ELS in military veterans. An important strength of this study is the use of repeated CV, where we show notable variance across 10 repeats of EPIC for each analysis. This is an important element of machine learning designs that should be considered in future studies, as it provides a reliability check that a feature selected as “important” is not a spurious result from one individual repeat of the classifier. We also report a specific function of self-reported depression and adult trauma exposure as important non-imaging markers that may distinguish people with PTSD and ELS in future data-driven designs. Further work will determine the generalizability of these findings in other cohorts with additional sources of clinical heterogeneity that are characteristic of PTSD populations.

## Acknowledgments and Disclosure

This work was supported by VISN6 MIRECC, VA Merit 1I01RX000389-01, NIH grants R01 NS086885, K23 MH073091, VA Merit 1I01CX000748, and NIH grants U54 EB020403 (BD2K), MH111671, and P41 EB015922. The authors have no conflicts of interest.

